# Genetic diversity, population structure and parentage analysis of Brazilian grapevine hybrids after half a century of genetic breeding

**DOI:** 10.1101/2022.07.30.502144

**Authors:** Geovani Luciano de Oliveira, Guilherme Francio Niederauer, Fernanda Ancelmo de Oliveira, Cinthia Souza Rodrigues, José Luiz Hernandes, Anete Pereira de Souza, Mara Fernandes Moura

## Abstract

In the 1940s, the Agronomic Institute of Campinas (IAC) started a grapevine breeding program to develop new cultivars adapted to the tropical and subtropical regions of Brazil. More than 2,000 crosses were carried out over 50 years, using 850 varieties as parents. However, among the thousands of hybrids developed by the program, only 130 are still maintained in the IAC grapevine germplasm collection. Little is known about their genetic makeup and usefulness for current breeding programs. In this study, we genotyped 130 Brazilian grape hybrids at 21 polymorphic microsatellite markers to evaluate the genetic diversity and population structure of the hybrids and verified their disclosed pedigrees. The results showed that the hybrid collection is highly diverse, with an expected heterozygosity (H_E_) of 0.80 and an observed heterozygosity (Ho) of 0.78. Strong structure in three subgroups based mainly on the usage and combination of parental groups was revealed by STRUCTURE software and confirmed by discriminant analysis of principal components (DAPC). Through molecular profiling analysis, fourteen synonyms, one homonym and one duplicate were identified. Parentage analysis confirmed 24 full parentages, as well as 33 half-kinships. In addition, 18 pedigrees were invalidated, and seven mislabeling events were identified. No compatible parent was identified for 33% of the IAC hybrids, highlighting severe genetic erosion in the IAC germplasm. The molecular characterization of the breeding hybrid bank collection contributes to our understanding of the genetic basis of the varieties, guiding the efficient utilization of available genetic diversity. Together, our results could be applied to other breeding programs and assist in the selection of parents, management of the breeding collection, and conservation of grapevine genetic resources.

## 1. Introduction

Brazil has different types of climatic and geographic conditions, which directly affect grapevine management strategies and production cycles (Pereira et al., 2020). The great socioeconomic importance of viticulture in Brazil and the considerable environmental variation in production zones located in temperate, subtropical, and tropical regions necessitates the development of different genetic breeding programs in the country with specific objectives in each region (Tecchio et al., 2018). In tropical and subtropical regions, traditional *Vitis vinifera* varieties often have adaptation problems in terms of bud dormancy, apical dominance, low fertility, and susceptibility to fungal diseases, factors that restrict production to a reduced number of varieties. Breeding programs in these regions of Brazil generally seek the development of new varieties through crossings of *Vitis vinifera* and *Vitis labrusca* varieties with interspecific hybrids and tropical wild species to combine the adaptation, productivity, resistance to diseases and quality of the grapes (Kok, 2014).

To promote the development of Brazilian tropical and subtropical viticulture, the Agronomic Institute of Campinas (IAC) started a grapevine breeding program in 1943, aiming to create new varieties that combine adaptation to tropical and subtropical environments, fruit quality according to purpose (table grapes, juice, and wine), productivity and disease resistance. In the beginning, a collection of European and American cultivars and French hybrids of the Seibel, Seyve Villard, and Couderc series was established. This material was evaluated, and the varieties that exhibited the best characteristics in terms of production, vine vigor, taste qualities, and resistance to biotic and abiotic factors were selected to be used as parents in the first set of crosses (De Santos Neto, 1971). To expand the genetic base, fourteen wild *Vitis* species, such as *Vitis rupestris, V. riparia, V. cinerea, V. caribaea, V. lincecumii*, and *V. labrusca*, were used because of their resistance to major pests and diseases. Based on the results of the first crosses in the IAC breeding program, hybrids with outstanding characteristics were used as parents (Pommer, 1993).

Over 50 years, since the beginning of the program, approximately 2,400 crosses have been performed using 850 different parental genotypes (Ferri and Pommer, 1995), leading to the release of varieties of wine grapes, table grapes, and rootstocks by the IAC. Among the thousands of hybrids developed by the program, only 130 are still maintained in the IAC grapevine germplasm; most of them were lost due to a variety of factors, such as resource limitations. The IAC hybrids are characterized by their complex pedigrees derived from crosses among three or more species, with a combination of alleles from different species of *Vitis*. Molecular characterization of these grapevine resources can help identify the genotypes that should be preserved and partially prevent or delay genetic erosion (Gago et al., 2022). Knowledge about the genetic diversity of germplasm resources is not only important for species protection but also necessary for the development and utilization of germplasm resources for crop improvement (Lassois et al., 2016).

The use of molecular markers has become an efficient method for genetic characterization and the determination of genetic relationships between germplasm accessions since the markers are not influenced by the environment and can be used in the early stages of plant development (Roychowdhury et al., 2014). Among the molecular markers identified in recent decades, microsatellites or simple sequence repeat (SSR) markers are highly polymorphic, abundant, reliably reproducible, relatively inexpensive to genotype, and transferable among several species of the genus *Vitis*, advantages that make them suitable and efficient for genetic analyses of grapevine resources (Cretazzo et al., 2022; This et al., 2004; Zarouri et al., 2015). In addition, SSRs provide unique fingerprints for cultivar identification. They are inherited following Mendelian codominant segregation, confirming their suitability for genetic resource characterization, genome mapping, assisted selection, and parentage analysis (Karastan et al., 2018; Khadivi et al., 2019; Mihaljević et al., 2020; Saifert et al., 2018; Vezzulli et al., 2019).

Historically, the putative parentage of new grape cultivars was recovered from breeders’ notes, which could be incomplete or inaccurate (Raimondi et al., 2017). Several studies have since used microsatellite markers to clarify the parentage relationships between grape cultivars, allowing for more accurate retrieval of breeding information by confirming or invalidating declared pedigrees and identifying new genetic relationships (Aliquó et al., 2017; Migliaro et al., 2019; Mihaljević et al., 2020).

Little is known about the genetic makeup of IAC grape hybrids and their usefulness for current breeding programs. Success in grapevine breeding depends on the understanding and use of the available gene pool of varieties and breeding clones (De Oliveira et al., 2020). Currently, there is great interest in understanding the genetic basis of complex traits and in discovering new germplasm traits that can be leveraged for efficient tropical grapevine breeding. IAC grape hybrids are thought to have high genetic value as a source of diversity. This valuable genetic resource can play an important role in the development of new varieties with favorable traits, such as adaptability to climate change, disease resistance, or an original flavor.

The goal of this study was to investigate, at the molecular level, grapevine hybrids developed over 50 years of breeding by the IAC Grapevine Breeding Program to assess their genetic diversity and population structure. Another aim was to clarify pedigree information to enable better categorization and advance understanding of the remaining interbreeds for use in future cross-breeding programs and the development of genetic conservation strategies.

## 2. Material and Methods

### 2.1. Plant material and DNA extraction

A total of 130 accessions of grapevine hybrids were analyzed in this study (Supplementary Table 1). The accessions were developed by the IAC Grapevine Breeding Program and belong to the Grapevine Germplasm Bank of the IAC located in Jundiaí, São Paulo (SP), Brazil. According to Köppen’s classification, the climate is of the Cfa type, that is, subtropical with dry winters and hot summers, and the soil in the area is classified as Cambisol Dystrophic Red (Santos et al., 2018). The vineyard was planted in October 2008 with a spacing of 2.5 m between rows and 1 m between vines (density of 4,000 vines per hectare). Vines were trained on a vertical shoot position (VSP) with a unilateral cord, and the wires were located at 1, 1.3, 1.5 and 1.8 m above the ground. Furthermore, some other cultural and phytosanitary management practices were also performed according to the standard practices for local growers in São Paulo (SP), Brazil. Each accession consisted of three clonally propagated plants pruned in August every year, leaving one or two buds per branch. After pruning, 2.5% hydrogen cyanamide was applied to induce and stimulate a more uniform budburst. For sampling, young leaves of a single plant were collected from each accession.

Total genomic DNA was extracted from young leaf material homogenized in a TissueLyser (Qiagen, Valencia, CA, USA) following the cetyltrimethylammonium bromide (CTAB) method previously described by Doyle (1991), with minor modifications to the extraction buffer. The buffer was composed of 2% CTAB, 700 mM NaCl, 1% 2-mercaptoethanol, 50 mM EDTA, 100 mM Tris-HCl pH 8.0, and 1% w/v polyvinylpyrrolidone (PVP). It was preheated, and the samples were incubated at 60°C in a water bath for 1 h, followed by two washes with chloroform-isoamyl alcohol (24:1), the addition of a 2/3 volume of cold isopropanol, and two washes with ethanol 70%. Finally, the pellet was dried and resuspended in 50 μl of water. The DNA concentration was quantified by using a NanoDrop 8000 (Thermo Scientific), and the quality was checked using 1% agarose gel electrophoresis.

### 2.2. Microsatellite analysis

A set of 21 microsatellites was selected to genotype the IAC hybrids in the study, including the set of nine SSR loci selected by the international scientific community for universal grapevine identification (Maul et al., 2012; This et al., 2004): VVS2 (Thomas and Scott, 1993), VVMD5, VVMD7 (Bowers et al., 1996), VVMD25, VVMD27, VVMD28, VVMD32 (Bowers et al., 1999), VrZAG62, and VrZAG79 (Sefc et al., 1999). Twelve additional markers previously developed to assess grapevine diversity were also included: VVIn74, VVIr09, VVIp25b, VVIn56, VVIn52, VVIq57, VVIp31, VVIp77, VVIv36, VVIr21, VVIu04 (Merdinoglu et al., 2005), and VrZAG64 (Sefc et al., 1999). Additional information about the loci is available in Supplementary Table 3.

Polymerase chain reaction (PCR) was performed using forward primers labeled with fluorescent dyes (6-FAM, PET, VIC, or NED) in a final volume of 10 μl containing 20 ng of template DNA, each primer at 0.2 μM, each dNTP at 0.2 mM, 2 mM MgCl_2_, 1× PCR buffer (20 mM Tris HCl [pH 8.4] and 50 mM KCl), and 1 U of Taq DNA polymerase. PCR amplifications were carried out using the following steps: 5 min of initial denaturation at 95°C followed by 35 cycles of 45 s at 94°C, 45 s at 58°C or 52°C (VVS2, VVMD7, VrZAG62 and VrZAG79), and 1 min 30 s at 72°C, with a final extension step of 7 min at 72°C. Capillary electrophoresis was conducted in an ABI 3500 machine (Applied Biosystems, Foster City, CA, USA). Allele calling was performed with Geneious software v. 8.1.9 (Kearse et al., 2012) using the internal GeneScan-600 (LIZ) Size Standard Kit (Applied Biosystems, Foster City, CA, USA).

### 2.3. Genetic diversity and population structure analysis

Descriptive statistics based on the genotyping data were generated using GenAlEx v. 6.5 (Peakall and Smouse, 2012) to indicate the number of alleles per locus (Na), effective number of alleles (Ne), observed heterozygosity (H_O_), expected heterozygosity (H_E_), and fixation index (F). The null allele frequency (r) and the polymorphism information content (PIC) were estimated using CERVUS 3.0.7 (Kalinowski et al., 2007). Discrimination power (D*j*) values were estimated to compare the efficiencies of microsatellite markers in varietal identification and differentiation (Tessier et al., 1999).

A model-based Bayesian analysis implemented in the software package STRUCTURE v. 2.3.4 (Pritchard et al., 2000) was used to determine the approximate number of genetic clusters (K) within the full dataset and to assign individuals to the most appropriate cluster. All simulations were performed using an admixture model, with 100,000 replicates treated as burn-in and 1,000,000 replicates for Markov chain Monte Carlo (MCMC) processes in ten independent runs. The number of clusters (K) tested ranged from 1 to 10.

Discriminant analysis of principal components (DAPC) as implemented in the R package adegenet was also performed using a nonparametric approach and free from Hardy–Weinberg constraints (Jombart et al., 2010). The *find.clusters* function was used to detect the number of clusters in the germplasm, running successive rounds of K-means clustering with increasing numbers of clusters (K). We used 20 as the maximum number of clusters. The optimal number of clusters was estimated using the Bayesian information criterion (BIC). The DAPC results are presented as multidimensional scaling plots.

### 2.4. Parentage and identity analysis

A search for compatible trio (parents and offspring) and duo (parent–offspring) combinations among the SSR profiles was carried out using a likelihood-based method implemented in CERVUS v.3.0.7 software (Kalinowski et al., 2007). The analysis was performed with molecular data from the IAC grapevine genetic database, including 280 additional genotypes (De Oliveira et al., 2020). Most of these accessions were European and American cultivars that were used as parents over the years by the IAC Grapevine Breeding Program.

The likelihood of each detected trio and duo was determined based on the natural logarithm of the overall likelihood ratio (LOD) score. The CERVUS program calculates allelic frequencies using a simulation approach and determines the confidence in parentage assignments by calculating critical values of the LOD score. One hundred thousand offspring were simulated with a proportion of 0.01 sampled parents, including the possibility of self-fertilization and the existence of relatives among potential parents. The maximum number of mismatching loci for trios and duos was set to 1, and the parentage relationship was considered significant when the trio or pair confidence was represented by a probability greater than 95%. Finally, the results of the analysis were compared with IAC historical records to verify declared parents.

To identify possible synonyms (different names for the same genotype), homonyms (common name for different genotypes) and duplicates, an individual identity analysis was also carried out using CERVUS software. The minimum number of matching loci was set to 13, and 1 fuzzy match was allowed.

## 3. RESULTS

### 3.1. Genetic diversity

One hundred thirty IAC grape hybrids were analyzed at 21 SSR loci, and a total of 258 alleles were detected (Table 1). The number of alleles per SSR locus (Na) ranged from 4 (VVIq57) to 21 (VVMD28), with an average of 12.28. The number of effective alleles per locus (Ne) varied from 2.12 (VVIq57) to 10.02 (VVMD28), with a mean value of 6.10.

**Table 1.** Genetic parameters of the 21 microsatellite loci obtained from 130 grapevine accessions. Na, number of alleles; Ne, number of effective alleles; H_O_, observed heterozygosity; H_E_, expected heterozygosity; PIC, polymorphism information content; D*j*, discrimination power; *r*, estimated frequency of null alleles. ^a^Standard error of mean values.

| Locus | $N_a$ | $N_e$ | $H_o$ | $H_E$ | PIC | $D_j$ | $r$ |
| --- | --- | --- | --- | --- | --- | --- | --- |
| VVIn74 | 12 | 5.70 | 0.75 | 0.82 | 0.80 | 0.83 | 0.04 |
| VVIr09 | 13 | 5.71 | 0.82 | 0.82 | 0.80 | 0.83 | -0.01 |
| VVIp25b | 11 | 3.15 | 0.60 | 0.68 | 0.64 | 0.69 | 0.05 |
| VVIn56 | 6 | 2.56 | 0.62 | 0.61 | 0.53 | 0.61 | -0.01 |
| VVIn52 | 11 | 7.89 | 0.71 | 0.87 | 0.86 | 0.88 | 0.10 |
| VVIq57 | 4 | 2.12 | 0.57 | 0.53 | 0.47 | 0.53 | -0.03 |
| VVip31 | 14 | 9.96 | 0.88 | 0.90 | 0.89 | 0.91 | 0.01 |
| VVip77 | 15 | 8.00 | 0.72 | 0.87 | 0.86 | 0.88 | 0.09 |
| VVIv36 | 10 | 4.45 | 0.79 | 0.78 | 0.74 | 0.78 | -0.01 |
| VVIr21 | 14 | 5.46 | 0.81 | 0.82 | 0.79 | 0.82 | 0.00 |
| VVIu04 | 14 | 5.88 | 0.81 | 0.83 | 0.81 | 0.83 | 0.00 |
| VVS2 | 14 | 7.05 | 0.85 | 0.86 | 0.84 | 0.86 | 0.00 |
| VVMD5 | 12 | 7.86 | 0.86 | 0.87 | 0.86 | 0.88 | 0.00 |
| VVMD7 | 14 | 9.41 | 0.92 | 0.89 | 0.88 | 0.90 | -0.01 |
| VVMD25 | 15 | 5.12 | 0.87 | 0.80 | 0.78 | 0.81 | -0.04 |
| VVMD27 | 13 | 6.38 | 0.85 | 0.84 | 0.83 | 0.85 | 0.00 |
| VVMD28 | 21 | 10.02 | 0.88 | 0.90 | 0.89 | 0.90 | 0.01 |
| VVMD32 | 12 | 4.50 | 0.64 | 0.78 | 0.75 | 0.78 | 0.09 |
| VrZAG62 | 12 | 7.10 | 0.93 | 0.86 | 0.85 | 0.86 | -0.04 |
| VrZAG79 | 12 | 6.50 | 0.88 | 0.85 | 0.83 | 0.85 | -0.02 |
| VrZAG64 | 9 | 3.18 | 0.65 | 0.68 | 0.65 | 0.69 | 0.03 |
| **Total** | 258 | 128 |  |  |  |  |  |
| **Mean** | 12.28 | 6.10 | 0.78 | 0.80 | 0.78 | 0.81 |  |
| **SE<sup>a</sup>** | 0.74 | 0.50 | 0.02 | 0.02 | 0.02 | 0.02 |  |

The mean observed heterozygosity (H_O_) and mean expected heterozygosity (H_E_) were very similar (0.78 and 0.80, respectively). A significantly high (>0.20) probability of null alleles (r) was not detected at any of the analyzed loci. The PIC had a mean value of 0.78, and the discrimination power (D*j*) was greater than 0.80 for 15 of the 21 loci, with a mean value of 0.81. When the PIC and D*j* of each locus were analyzed together, 13 loci exhibited high values for both indexes (>0.80).

Of the 258 SSR alleles found, 48.1% displayed a frequency greater than 5% and were classified as common alleles, 35.6% had frequencies between 1% and 5% and were considered less common alleles, and 16.3% had a frequency less than 1% and were rare alleles, suggesting that this collection, despite originating from the same breeding program, includes great biodiversity.

### 3.2. Population structure analysis

The STRUCTURE analysis indicated relatedness among the 130 accessions, with the highest ΔK value for K = 3, suggesting that three genetic clusters were sufficient to interpret our data (Supplementary Fig. 1). Based on a membership probability threshold of 0.70, 22 hybrids were assigned to cluster SV (Seibel hybrids x *V. vinifera*), 41 hybrids were assigned to cluster MG (Muscat table grape offspring), and 44 hybrids were assigned to cluster TV (tropical vine offspring). Twenty-three hybrids were not assigned to defined clusters and were assigned to the admixed group (Fig. 1).

**Fig. 1.**
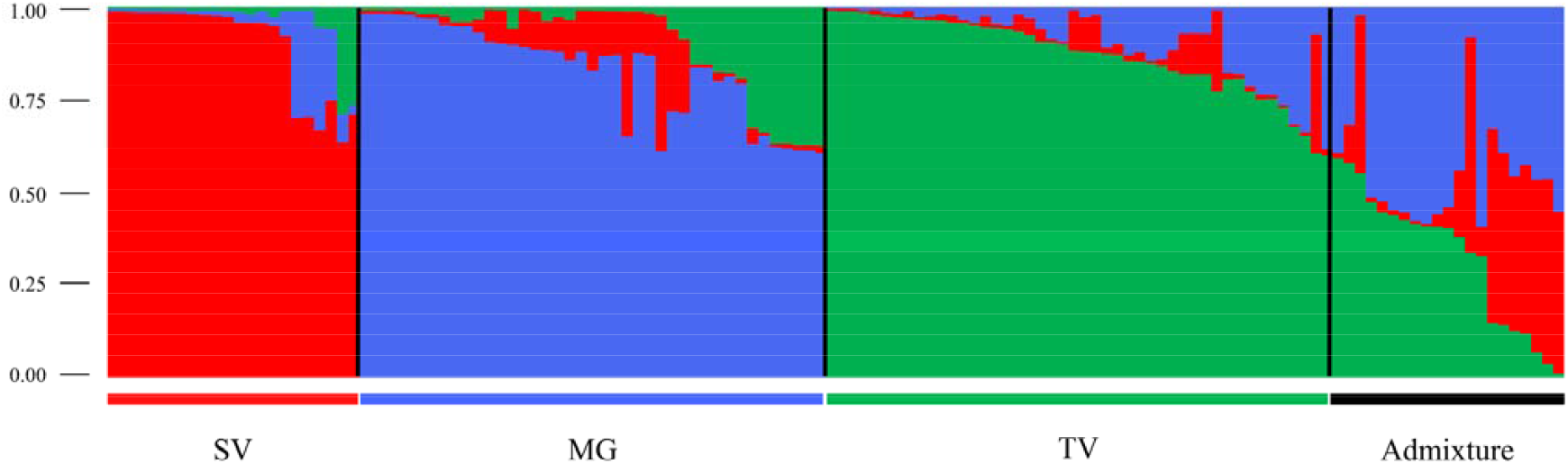
Bar graph of the estimated membership coefficient (q) for each of the 130 individuals. Each genotype is represented by a single vertical line, which is partitioned into colored segments in proportion to the estimated membership in each cluster. Cluster SV: predominantly wine hybrids; cluster MG: predominantly table grape hybrids; cluster TV: hybrids originating from crosses with wild *Vitis* (tropical vines); Admixture: interspecific hybrids with a membership of q < 0.70.

The clustering level was mainly based on the use and combination of parental gene pools. Cluster SV was formed primarily by wine hybrids originating from crosses between Seibel hybrids and wine grape varieties of *V. vinifera*. Cluster MG was composed of table grape hybrids originating from crosses with Muscat grapes. In the TV cluster, there was no clear discrimination based on human use, and hybrids for wine, table grapes and rootstock were found in this group. However, all these hybrids were developed from crosses with tropical vines (wild *Vitis*). The hybrids in the admixed group exhibited more complex origins, and some of them had associations with all three clusters.

Additionally, DAPC was performed with no prior information about the groupings of the evaluated accessions. Inspection of the BIC values revealed that the division of the accessions into three clusters was the most likely scheme to explain the variance in this set of genotypes (Supplementary Fig. 2). In the preliminary step of data transformation, the maintenance of 90 principal components (PCs) allowed DAPC to explain 99% of the total genetic variation. The DAPC scatter plot, based on the first and second discriminant functions, showed the distribution of the three groups (Fig. 2), revealing great genetic differentiation between them and low variance within them.

**Fig. 2.**
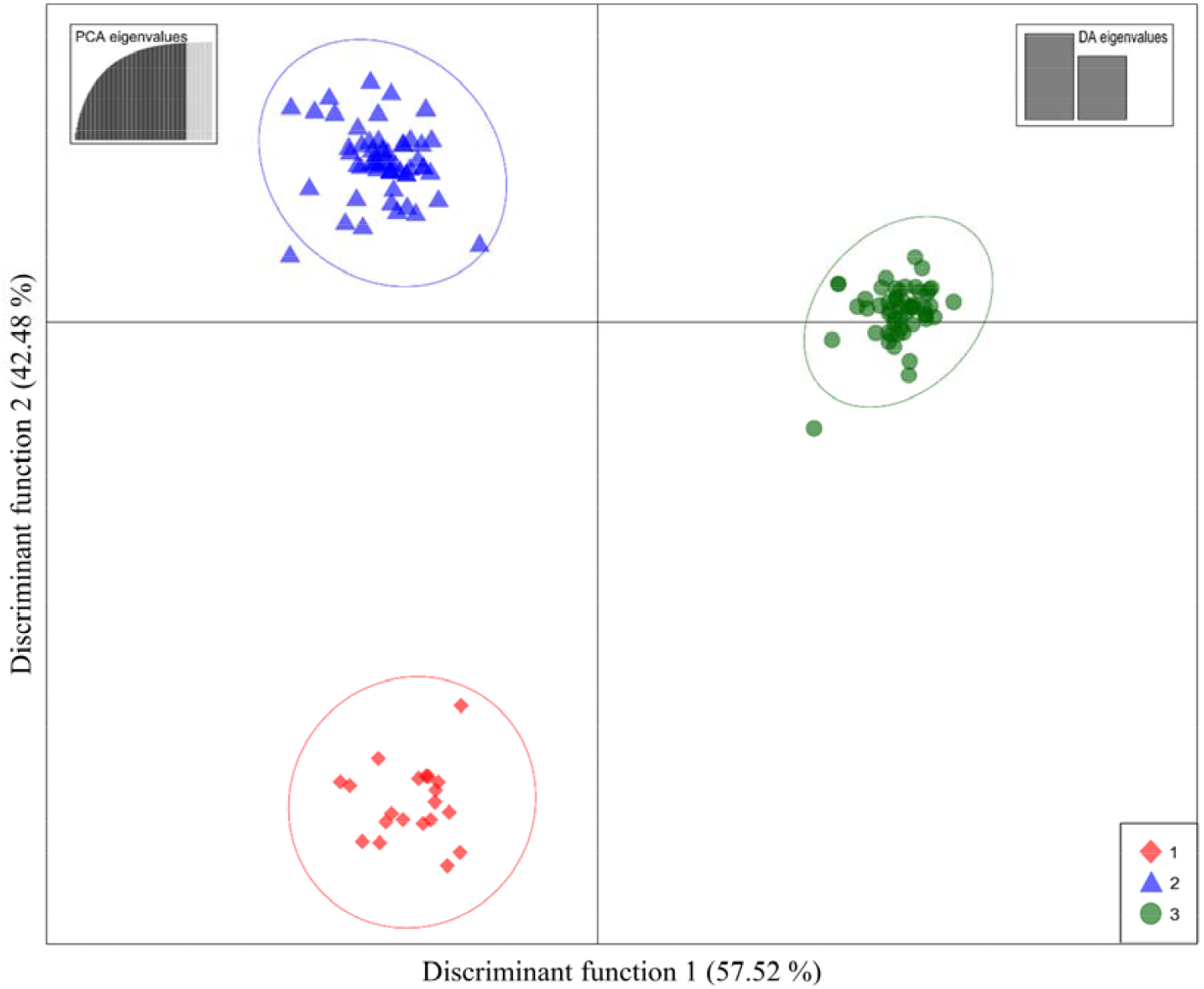
DAPC scatterplots based on the K-means algorithm used to identify the best number of clusters. Dots represent individuals, and the clusters are presented in different colors. The individuals were allocated to three clusters: 1 (red), wine hybrids obtained by crossing Seibel hybrids with *V. vinifera* wine cultivars; 2 (blue), table grape hybrids obtained through crosses with fine Muscat grapes; and 3 (green), hybrids obtained through crosses with wild *Vitis*.

The allocation of individuals to clusters according to the DAPC results showed several similarities to that resulting from STRUCTURE, and both analyses showed the same pattern of clustering. Clusters 1 (red lozenge), 2 (blue triangle), and 3 (green circle) from the DAPC reflected the subgroups SV, MG, and TV detected by STRUCTURE, respectively. According to the DAPC grouping results, 20 hybrids were allocated to Cluster 1, 57 to Cluster 2, and 53 to Cluster 3. Cluster 1 and subgroup SV showed high grouping similarity, with all individuals from Cluster 1 present in subgroup SV. Regarding the similarity between the other groupings, 32 hybrids (78%) from the MG subgroup were found in Cluster 2, and 33 hybrids (75%) from the TV subgroup were allocated to Cluster 3. Individuals in the admixed group from STRUCTURE analysis were basically distributed in DAPC Clusters 2 and 3, with 14 hybrids (61%) assigned to Cluster 2 and eight (35%) to Cluster 3. DAPC analysis proved to be more discriminating, and no cases of overlap between clusters were observed, indicating more delineated genetic structure.

#### 3.2.1 Parentage analysis

To improve the search for possible first-order kinship relationships, the genetic profiles of 280 accessions stored in the IAC grapevine genetic database (De Oliveira et al., 2020) were added to parentage analyses, resulting in a total of approximately 410 genotypes. The critical LOD values determined by simulation for strict confidence (95%) of parentage were 19.00 and 6.40 for trios (offspring and two inferred parents) and duos (parent–offspring), respectively. Offspring resulting from self-pollination were not detected. A total of 33 compatible trios were identified with a high confidence level using a maximum of one mismatched locus as the threshold, with LOD values ranging from 27.78 to 57.96 (Table 2). The complete pedigrees of 24 IAC hybrids reported in the IAC records were validated, while for three identified trios, one parent was validated and the other was not (Supplementary Table 1). For nine IAC hybrids, both declared parents were invalidated, and for two of them, no other possible parent (consistent with the offspring’s SSR profiles) was found in the IAC grapevine genetic database.

**Table 2.** Putative full parentages of 33 IAC grapevine hybrids inferred based on the maximum likelihood approach. * A maximum of only one locus mismatch was allowed, and the parentage relationship was considered significant when the trio confidence probability was greater than 95% (LOD ≥ 19.00).

| Offspring<br>ID number | Offspring name | First candidate parent | Second candidate<br>parent | Trio loci<br>compared | Trio loci<br>mismatches | Trio LOD<br>score* |
| --- | --- | --- | --- | --- | --- | --- |
| 1 | IAC 16-02 | Ravat 34 | Muscat Hamburg | 20 | 0 | 29.75 |
| 2 | IAC 21-14 (Madalena) | Ravat 34 | Moscato Giallo | 19 | 1 | 27.78 |
| 3 | IAC 23-08 | Ravat 34 | Muscat Hamburg | 20 | 1 | 32.09 |
| 5 | IAC 74-01 (Iara) | Seibel 10096 | Syrah | 19 | 0 | 38.32 |
| 6 | IAC 74-02 | Seibel 10096 | Syrah | 21 | 0 | 36.44 |
| 8 | IAC 116-31 (Rainha) | Seibel 7053 | Pinot Noir | 21 | 0 | 40.25 |
| 13 | IAC 192-54 | Seibel 8712 | Muscat Hamburg | 21 | 0 | 37.48 |
| 21 | IAC 313 (Tropical) | Golia | *Vitis cinerea* | 20 | 1 | 57.96 |
| 24 | IAC 339-03 | Muscat Hamburg | *Vitis cinerea* | 20 | 0 | 39.43 |
| 25 | IAC 341-02 | Moscatel Rosado | *Vitis cinerea* | 20 | 0 | 47.32 |
| 29 | IAC 388 (Santa Tereza) | Italia | IAC 82-01 | 21 | 0 | 35.55 |
| 33 | IAC 405-06 | Moscatel Rosado | *Vitis cinerea* | 21 | 0 | 46.84 |
| 36 | IAC 457-11 (Iracema) | Niagara Branca | Sultanina | 21 | 1 | 39.25 |
| 37 | IAC 460-01 | Highland | Sultanina | 18 | 0 | 40.36 |
| 40 | IAC 496-15 | Seibel 7053 | Gewurztraminer | 20 | 0 | 45.73 |
| 71 | IAC 775-26 (Aurora) | Niagara Branca | Sultanina | 21 | 0 | 39.25 |
| 80 | IAC 871-05 (Geni) | IAC 501-06 (Soraya) | IAC 544-14 | 21 | 0 | 38.87 |
| 81 | IAC 871-13 (A Dona) | IAC 501-06 (Soraya) | IAC 544-14 | 21 | 0 | 38.87 |
| 82 | IAC 871-18 | IAC 501-06 (Soraya) | IAC 544-14 | 20 | 0 | 42.15 |
| 83 | IAC 871-41 (Patricia) | IAC 501-06 (Soraya) | IAC 544-14 | 21 | 0 | 41.21 |
| 99 | IAC 966-01 | Seibel 7053 | Pinot Noir | 21 | 0 | 40.25 |
| 114 | IAC 1742 | Muscat Hamburg | Niagara Branca | 19 | 0 | 32.88 |
| 119 | IAC Juliana | IAC 21-14 (Madalena) | Italia | 21 | 0 | 33.36 |
| 121 | SR 496-09 | Seibel 7053 | Gewurztraminer | 20 | 0 | 47.96 |
| 122 | SR 496-15 (Dr. Júlio) | Seibel 7053 | Gewurztraminer | 19 | 0 | 44.70 |
| 123 | SR 496-25 | Seibel 7053 | Gewurztraminer | 21 | 1 | 42.08 |
| 124 | SR 5010-08 | Seibel 7053 | Seibel 10096 | 21 | 1 | 37.21 |
| 125 | SR 5010-21 | Seibel 7053 | Seibel 10096 | 20 | 1 | 37.21 |
| 126 | SR 501-17 (IAC Ribas) | Seibel 7053 | Syrah | 21 | 0 | 46.83 |
| 127 | SR 5012-34 (Dona Emília) | Seibel 7053 | Cabernet Sauvignon | 20 | 1 | 31.84 |
| 128 | SR 501-33 | Seibel 7053 | Syrah | 21 | 1 | 39.75 |
| 129 | SR 507-38 | Seibel 7053 | Sémillon | 19 | 1 | 28.62 |
| 130 | SR 507-08 | Seibel 7053 | Cabernet Sauvignon | 21 | 1 | 34.83 |

A total of 42 compatible duos were also identified, and all of them were recognized as cases of putative direct (first-degree) relationships (Supplementary Table 4). The partial pedigrees of 33 IAC hybrids reported in the IAC records were validated, while for nine IAC hybrids, the identified parent did not correspond to any of the declared parents. Moreover, no reliable trios or duos within the IAC grapevine genetic database were found for the other 43 genotypes. The two most common varieties that emerged as a parent in 18 proposed trios and five duos were ‘Seibel 7053’ (syn. ‘Chancellor’) and ‘Muscat Hamburg’. The next most recurrent parent in seven crosses (four trios and three duos) was the hybrid IAC 544-14, which had an unverifiable pedigree, as its declared parents were IAC hybrids that are now extinct.

#### 3.2.2 Genotype identity

Among the 130 IAC hybrids analyzed, 14 synonyms were identified, with hybrids having the same molecular profile but identified with different names (Supplementary Table 5). In addition, one case of duplication and one case of homonymy were detected. The two hybrids labeled IAC 746-03 showed the same molecular profile, while the molecular profiles of the two hybrids labeled IAC 514-6 were different.

Through pedigree validation, seven synonyms were identified as possibly mislabeled, and their correct identification is proposed in Supplementary Table 5. It is possible that the other seven synonyms identified in this study were also due to mislabeling; however, it was not possible to propose a correct identity in these cases because either no parent was identified in the parentage analysis or both hybrids of the synonym group had the same pedigree.

## 4. DISCUSSION

### 4.1. Genetic diversity

The results of this study revealed high levels of heterozygosity among the evaluated genotypes, with a high percentage of less common and rare alleles (52%). Since heterozygosity is an indicator of genetic variability in a population and is related to the polymorphic nature of each locus, these results highlight the potential of this genetic material as a source of genetic diversity. The wide abundance of parents used in the crosses and the different purposes of the breeding program were probably the factors responsible for the high genetic diversity we observed. Among the 850 genotypes used as parents by the IAC breeding program, there were *V. vinifera* cultivars from different countries, wild species and intra- and interspecific hybrids. Approximately 2,400 combinations of these parents were bred to obtain wine, table and rootstock grape varieties adapted to conditions in Brazil (Ferri and Pommer, 1995).

We detected an H_E_ of 0.80 across the entire hybrid set at the 21 evaluated loci (Table 1). This result is similar to those found in other grapevine collections characterized by an abundance of interspecific hybrids (Migliaro et al., 2019; Schuck et al., 2009) but greater than that of collections composed only of *V. vinifera* accessions (De Lorenzis et al., 2014; Riaz et al., 2018). Laucou et al. (2011) and Emanuelli et al. (2013) showed that the genetic diversity found in non-*vinifera* varieties was higher than that in the *V. vinifera* sector, indicating that taxonomically broader genotypes contribute to an increase in genetic diversity, as expected by the heterogeneity of IAC hybrids, since most have wild *Vitis* in their genealogy.

The large number of alleles obtained by the 21 SSR primer set positively impacted the PIC and discrimination power (D*j*). No locus was identified with a high frequency of null alleles (> 0.20). According to the classification of Botstein et al. (1980), all the loci in the study can be considered highly informative (PIC > 0.50), except for VVIq57 (0.47). This locus also presented the lowest D*j* value, certainly due to its reduced number of alleles (4), which limits the power to distinguish genotypes. All 20 other SSR loci analyzed proved to be adequate for grape cultivar discrimination and can be considered an efficient set for genetic diversity studies.

### 4.2. Cluster analysis and genetic structure

Genetic structure was impacted by the different objectives and strategies adopted by the IAC breeding program, such as the development of grape varieties for wine, table grapes and rootstock adapted to the climatic conditions in Brazil through crosses between *V. viniferas* cultivars, complex hybrids and wild *Vitis* species known as tropical vines. Population structure analysis using STRUCTURE software revealed the presence of three primary clusters in our set of hybrids (Fig. 1), two of which were strongly based on human usage and the other of which had no clear distinction regarding use but was strongly associated with tropical vines.

Most of the hybrids developed for use as wine grapes were concentrated in the SV cluster. Based on the analysis of the genealogy of the hybrids of this cluster, there was a clear direction in the use of Seibel series hybrids crossed with wine grape cultivars of *V. vinifera*. Seibel series hybrids were widely used in the state of São Paulo from the 1930s through the 1950s, and they exhibited good productivity, good affinity with the rootstocks used in the region, and satisfactory resistance to the main pests and diseases. However, they had some problems regarding the quality of the wine produced (Souza and Martins, 2002). On the other hand, the *V. vinifera* cultivars known for producing high-quality wines exhibited weak adaptation to the climatic conditions in Brazil. The SV cluster reflects one of the strategies used in the breeding program to develop cultivars capable of producing high-quality wines under the tropical and subtropical conditions in Brazil. Basically, all hybrids in this cluster were obtained from crosses of the Seibel series with *V. vinifera* cultivars, except SR 5010-08 and SR 5010-21, which were obtained by crossing two Seibel hybrids.

The MG cluster was formed by table grape hybrids with a predominance of genealogies based on crosses with Muscat grapes. In the 1950s, there was a high market demand for Muscat-flavored table grapes in Brazil, for which high prices were paid. Most of the Muscat grapes used in the country had adaptability problems, such as cluster rot, berry splitting, and susceptibility to fungal diseases, mainly downy mildew and powdery mildew. Given this scenario, one of the focuses of the breeding program was to obtain new varieties resistant to the main fungal diseases, with satisfactory development under the conditions in Brazil, and with fruits of high palatability, high sugar content, low acidity, and muscatel flavor (Neto and Almeida, 1955). Most of the hybrids in the MG cluster were the result of this approach, arising mainly from crosses with ‘Moscatel Branco’ (‘Moscato Giallo’), ‘Moscatel Rosado’, ‘Muscat Hamburg’, and ‘Italia’.

Unlike previous clusters, there was no clear discrimination based on usage in the TV cluster, and hybrids for wine, table grapes, and rootstock were found in this group. However, all hybrids have wild *Vitis* in their genealogy in common. Tropical vines were used intensively in the IAC breeding program to promote climate adaptability and disease resistance. These vines have small fruits with a low percentage of pulp, and their chemical composition lack a satisfactory balance, not meeting the requirements for table or wine grapes. However, because of their characteristics related to vigor, resistance, productivity, and adaptation to regions with high humidity and temperature during the summer, these species played an important role in efforts to expand the genetic base of the new Brazilian varieties (Ferri and Pommer, 1995).

A small number of the hybrids remained admixed, with evidence of a greater genetic complexity of these genotypes. The intra- and interspecific crossings carried out during breeding cycles in search of novelties and hybrid vigor promoted the miscegenation of grapevine cultivars, resulting in hybrids with a heterogeneous genetic composition (De Oliveira et al., 2020). The admixed group hybrids certainly carry alleles from different gene pools; they occupy an intermediate position and belong to more than one cluster. The hybrids IAC 339-03 and IAC 393-04 are examples of this condition. These hybrids resulted from crosses between the cultivar ‘Muscat Hamburg’ and the tropical vines *V. smalliana* and *V. shuttleworthii* x *V. rufotomentosa*, respectively. The mixture of gene pools was detected by STRUCTURE, which assigned a membership probability threshold of approximately 0.5 to the MG and TV clusters, representing the genetic clusters of the two parental cultivars (Muscat grapes and tropical vines). The other hybrids from the Admixture group exhibited a similar or even more complex origin than these examples, and some of them had associations with the three clusters simultaneously.

Individuals in the STRUCTURE admixed group were predominantly distributed between DAPC clusters 2 and 3, possibly because the DAPC analysis minimizes within-group genetic variance and maximizes between-group genetic variance. The genotypes in this study are the result of human manipulation of cultivars (displacements, breeding, and clonal propagation); therefore, deviations from Hardy–Weinberg equilibrium (HWE) are expected. This feature can lead to greater accuracy in the DAPC results since this method does not assume the absence of linkage disequilibrium or specific models of molecular evolution to identify genetic clusters (Jombart et al., 2010). The use of more than one clustering method prevents erroneous inferences from being adopted in the allocation of genotypes within a given subgroup, ensuring that the obtained result is not an artifact of the technique used (De Oliveira et al., 2020). The different grouping criteria of the two analyses reflect some attribution differences, mainly between the subgroups of STRUCTURE MG and TV with DAPC Clusters 2 and 3, probably because both genetic clusters include hybrids with a genetic basis related to wild *Vitis*. However, the similarity between the two analyses was greater than 75%, resulting in the same grouping pattern. In addition, the results of the two analyses can be used in a complementary way, since the grouping by DAPC provides greater distinction between the genotypes and the analysis by STRUCTURE adds information regarding the genetic composition of the hybrids, providing information about the proportion of each cluster in their genome.

The clustering of individuals provides interesting clues for increasing diversity in breeding programs and germplasm collections (Pandey et al., 2021). Knowledge about the genetic structure of IAC hybrids will certainly help minimize the use of closely related genotypes as parents in breeding programs, avoiding the risk of inbreeding depression and the reduction of genetic variation. Information regarding genetic diversity, population structure, and molecular markers may facilitate the selection of desirable traits in grapes and is important for ensuring the conservation of genetic resources.

### 4.3. Parentage analysis and its use in determining genotype identity

Among the IAC hybrids analyzed in this study, only 30 (23.07%) were actually released as varieties. The others remained exclusively in the IAC grapevine germplasm without any published genealogy information. In this study, we made available the genealogy information of all 130 hybrids (Supplementary Table 1) recovered through research carried out in the breeder’s notes and institution’s internal records. We also performed parentage analysis of these hybrids with molecular data for the first time and used the results to validate the parentage declared in the historical records.

Nine trios and eight duos had their declared pedigrees invalided by parentage analysis. In all IAC hybrids examined, ‘Seibel 11342’ was invalidated as a parent, with ‘Ravat 34’ being the true parent. The correct identification of this cultivar in the IAC germplasm was suggested in a recent study (De Oliveira et al., 2020) and was confirmed in this study as a mislabeling that likely occurred beginning with the first crosses in 1944, indicating that ‘Seibel 11342’ was not introduced in the IAC grapevine breeding program. The use of ‘Ravat 34’ instead of ‘Seibel 11342’ in a substantial number of crosses increased the inaccuracy of the breeder’s data, since important hybrids such as IAC 21-14 Madalena and IAC 138-22 Máximo were released with incorrect genealogy information and were later used as parents in new crosses.

Some hybrids with invalidated pedigrees were identified as synonyms by identity analysis (Supplementary Table 5). Since most of the IAC hybrids were kept exclusively in the IAC germplasm, the synonyms found were probably “internal synonyms”, originating from cases of misnaming that occurred over the years. Misidentification in breeding programs is common, especially for ancient clonal species such as *Vitis* spp., and it can occur during material propagation, during the planting and duplication of collections, or even during seedling selection (Raimondi et al., 2017). Through pedigree validation, we proposed the correct identification of seven synonyms. In these cases, the synonym presented the same genetic profile as a hybrid with a validated pedigree. For the other seven synonyms, correct identification was more complex, since some had extinct parents and others had the same parents. Basic ampelographic data showed that the 14 synonyms found share the same phenotype (Supplementary Table 6), reinforcing the results detected by molecular analysis. However, more ampelographic and passport data are necessary for these synonyms to check for true synonym status (not yet known), to identify possible somatic mutations not detected with a small number of SSR markers (Cipriani et al., 2010; Liang et al., 2015) and to discard cases of false synonymy resulting from grafting errors or erroneous former morphological identification (Lassois et al., 2016; Maul and Töpfer, 2015).

In addition to these synonyms, one case of homonymy was also found. The hybrids IAC 514-6 (ID: 46) and IAC 514-6 (Maria) (ID: 47) shared the same name but not the same genetic profile. In the literature, this variety has been described as a seedless white table grape (Pommer, 1993; Pommer et al., 1995), and according to IAC phenotyping data (unpublished), only the IAC 514-6 (Maria) (ID: 47) genotype matches these descriptions. Both genotypes corresponded to white table grapes, but only IAC 514-6 (Maria) (ID: 47) was a seedless grape, and the other presented well-developed seeds. This evidence points to the hybrid IAC 514-6 (Maria) (ID: 47) as the correct variety. IAC 514-6 (ID: 46) was another genotype that could not be identified, likely another result of mislabeling.

In this study, no compatible parent was identified for 43 IAC hybrids (33%) within the IAC grapevine genetic database, and for another 42 (32.3%), only one compatible parent was detected. The low number of reconstructed trios (both parents and offspring) points to the severe genetic erosion of the IAC germplasm since the late 1980s. Most hybrids with unverifiable parents were the result of crosses between genotypes developed by the IAC breeding program that went extinct. At the beginning of the breeding program, a large number of crosses were carried out, and numerous hybrids were obtained. Many of these hybrids were not released as cultivars but played an important role as intermediaries in the use of wild species, often being used as parents (Ferri and Pommer, 1995). The importance and justification for the preservation of this large volume of local genotypes were overlooked, since most of them were not economically interesting at the time; the lack of financial support resulted in the loss of a large part of the IAC genetic resources.

Declared pedigrees are not necessarily a reliable tool, either because they are often too generic (such as *V. shuttleworthii* x *V. rufotomentosa*) or because the declared parents do not match the true parents due to mislabeling issues (Migliaro et al., 2019). Therefore, genetic data analysis is essential to verify the consistency of declared parents, and it can help correct cases of mislabeling and ensure true variety identification (Raimondi et al., 2017). Microsatellite markers are among the most commonly used molecular markers for genetic analysis in grapevines since the alleles are inherited via Mendelian codominant segregation, confirming their suitability for investigating hereditability and cultivar parentage (Aliquó et al., 2017; De Lorenzis et al., 2014; Mihaljević et al., 2020; Sefc et al., 2009). In this study, the 21 SSRs used were valuable for drawing robust conclusions regarding first-degree relationships, supporting or questioning known information, suggesting new possible parentage, and identifying probable cases of misidentification. This number of markers has been frequently used in parental analysis studies, allowing efficient access to information on the ancestry of grapevine cultivars in regard to a polymorphic set that is well distributed throughout the genome (Aliquó et al., 2017; Lacombe et al., 2013; Mihaljević et al., 2020).

## 5. CONCLUSIONS

Despite the serious genetic erosion that occurred in the IAC grapevine germplasm, this study revealed that there is still a high level of genetic diversity present in the set of conserved hybrids developed by the breeding program. However, this loss of genetic resources made it impossible to fully validate the pedigrees of most individuals, since many IAC hybrids used as key parents were no longer present in the collection.

The combined results from the parentage and identity analyses allowed us to identify cases of genotype mislabeling, information that is extremely useful for curating the collection. Additional phenotypic and passport data checking is necessary to address pending identification questions. The overall diversity structure was shown to be rather strong and coincided with the usage of the varieties and the strategies adopted by the breeding program based on combinations of parental groups.

Many of the hybrids in this study were not properly recognized as cultivars and can be considered a source of genetic diversity with the potential for utilization; they could be used to obtain new varieties that may exhibit crucial features for developing sustainable viticulture in tropical and subtropical areas. All these data point to the importance and justification of preserving these genotypes in germplasm repositories.

## Acknowledgements

This study was funded by the São Paulo Research Foundation (FAPESP) grants 2018/13539-9 and 2020/12938-7 and Conselho Nacional de Desenvolvimento Científico e Tecnológico (CNPq) grant 404041/2021-3. Coordination for the Improvement of Higher Education Personnel (CAPES) provided fellowships to GLO (grant 88887.479802/2020-00), GFN (grant 88887.513247/2020-00), and CSR (grant 88882.317454/2019-01). FAO received a PostDoctoral fellowship from the São Paulo Research Foundation (FAPESP) (grant 2018/18527–9). The funders had no role in the study design, data collection and analysis, decision to publish, or preparation of the manuscript.

## CRediT authorship contribution statement

**Geovani Luciano de Oliveira:** Conceptualization, Data curation, Methodology, Formal analysis, Visualization, Investigation, Writing - original draft; Writing - review & editing. **Guilherme Francio Niederauer:** Methodology, Writing - original draft. **Fernanda Ancelmo de Oliveira:** Conceptualization, Visualization, Writing - review & editing. **Cinthia Souza Rodrigues:** Writing - review & editing, Investigation. **José Luiz Hernandes:** Methodology, Writing - review & editing. **Anete Pereira de Souza:** Funding acquisition, Project administration, Resources, Supervision. **Mara Fernandes Moura:** Conceptualization, Visualization, Writing - review & editing, Funding acquisition, Project administration, Resources, Supervision.

