## Supplementary file 2 for "Genetic diversity, population structure and parentage analysis of Brazilian grapevine hybrids after half a century of genetic breeding"

**Supplementary Table 3.** Microsatellite markers used in this study.

| **SSR Locus Name** | **Linkage Group** | **Microsatellite Repeat Motif** | **Sequence** | **Reference** |
| --- | --- | --- | --- | --- |
| VVIn74 | 19 | (AG)_n_ | F: TTGGTTGAGGGAAAAAGGAAAG | Merdinoglu et al. (2005) |
|  |  |  | R: AAGAAATATGAGGTTGGGTGAG |  |
| VVIr09 | 18 | (GA)_n_ | F: AAGTGTGTTTGACTCCAGAAAA | Merdinoglu et al. (2005) |
|  |  |  | R: ACTGATCAAACTTCTCTAGAGA |  |
| VVIp25b | 4 | (GA)_n_ | F: AAGAAAAGAAAAAAGGTGGGGG | Merdinoglu et al. (2005) |
|  |  |  | R: CCAGAGTCTCTTGTCATTCATC |  |
| VVIn56 | 7 | (AC)_n_ | F: GCCAAGTAGCCAAATTATAGAA | Merdinoglu et al. (2005) |
|  |  |  | R: TTTATGCTCCGTGGTTTGAAAT |  |
| VVIn52 | 16 | (GA)_n_ | F: TTTTTGTCGACAACCAAGAAGG | Merdinoglu et al. (2005) |
|  |  |  | R: CCATACACCTCACTAAATTCAG |  |
| VVIq57 | 1 | (GT)_n_ | F: TGAACCTCTATCTTCCATGTAG | Merdinoglu et al. (2005) |
|  |  |  | R: TATGCCCTGTTACTATCTATGA |  |
| VVIp31 | 19 | (GA)_n_ | F: TATCCAAGAGACAAATTCCCAC | Merdinoglu et al. (2005) |
|  |  |  | R: TTCTCTTGTTTCCTGCAAATGG |  |
| VVIp77 | 4 | (CT)_n_ | F: GGGTACGAATTCTCATGTTATC | Merdinoglu et al. (2005) |
|  |  |  | R: CCATCTTCAATGAATCAATCCC |  |
| VVIv36 | 7 | (GA)_n_ | F: AGCAAGAGCTCAATCCATAAAA | Merdinoglu et al. (2005) |
|  |  |  | R: TAGTTAGTTGGCATAGTAGGTT |  |
| VVIr21 | 10 | (GA)_n_ | F: TTTCCCTTCTCACTCAATGATG | Merdinoglu et al. (2005) |
|  |  |  | R: GAAAAGAAAACAGTGTATCCCC |  |
| VVIu04 | 18 | (TC)_n_ | F: ACAAAAGCGGAAACGATCGAAT | Merdinoglu et al. (2005) |
|  |  |  | R: AGAAGACCTATTTTTCCTGTGG |  |
| VVS2 | 11 | (GA)_n_ | F: CAGCCCGTAAATGTATCCATC | Thomas and Scott (1993) |
|  |  |  | R: AAATTCAAAATTCTAATTCAACTGG |  |
| VVMD5 | 16 | (CT)_n_AT(CT)_n_ATAG(AT)_n_ | F: CTAGAGCTCGCCAATCCAA | Bowers et al. (1996) |
|  |  |  | R: TATACCAAAAATCATATTCCTAAA |  |
| VVMD7 | 7 | (CT)_n_ | F: AGAGTTGCGGAGAACAGGAT | Bowers et al. (1996) |
|  |  |  | R: CGAACCTTCACACGCTTGAT |  |
| VVMD25 | 11 | (CT)_n_ | F: TTCCGTTAAAGCAAAAGAAAAAGG | Bowers et al. (1999) |
|  |  |  | R: TTGGATTTGAAATTTATTGAGGGG |  |
| VVMD27 | 5 | (CT)_n_ | F: GTACCAGATCTGAATACATCCGTAAGT | Bowers et al. (1999) |
|  |  |  | R: ACGGGTATAGAGCAAACGGTGT |  |
| VVMD28 | 3 | (CT)_n_ | F: AACAATTCAATGAAAAGAGAGAGAGAGA | Bowers et al. (1999) |
|  |  |  | R: TCATCAATTTCGTATCTCTATTTGCTG |  |
| VVMD32 | 4 | (CT)_n_ | F: TATGATTTTTTAGGGGGGTGAGG | Bowers et al. (1999) |
|  |  |  | R: GGAAAGATGGGATGACTCGC |  |
| VrZAG62 | 7 | (GA)_n_ | F: GGTGAAATGGGCACCGAACACACGC | Sefc et al. (1999) |
|  |  |  | R: CCATGTCTCTCCTCAGCTTCTCAGC |  |
| VrZAG79 | 5 | (GA)_n_ | F: AGATTGTGGAGGAGGGAACAAACCG | Sefc et al. (1999) |
|  |  |  | R: TGCCCCCATTTTCAAACTCCCTTCC |  |
| VrZAG64 | 10 | (GA)_n_ | F: TATGAAAGAAACCCAACGCGGCACG | Sefc et al. (1999) |
|  |  |  | R: TGCAATGTGGTCAGCCTTTGATGGG |  |

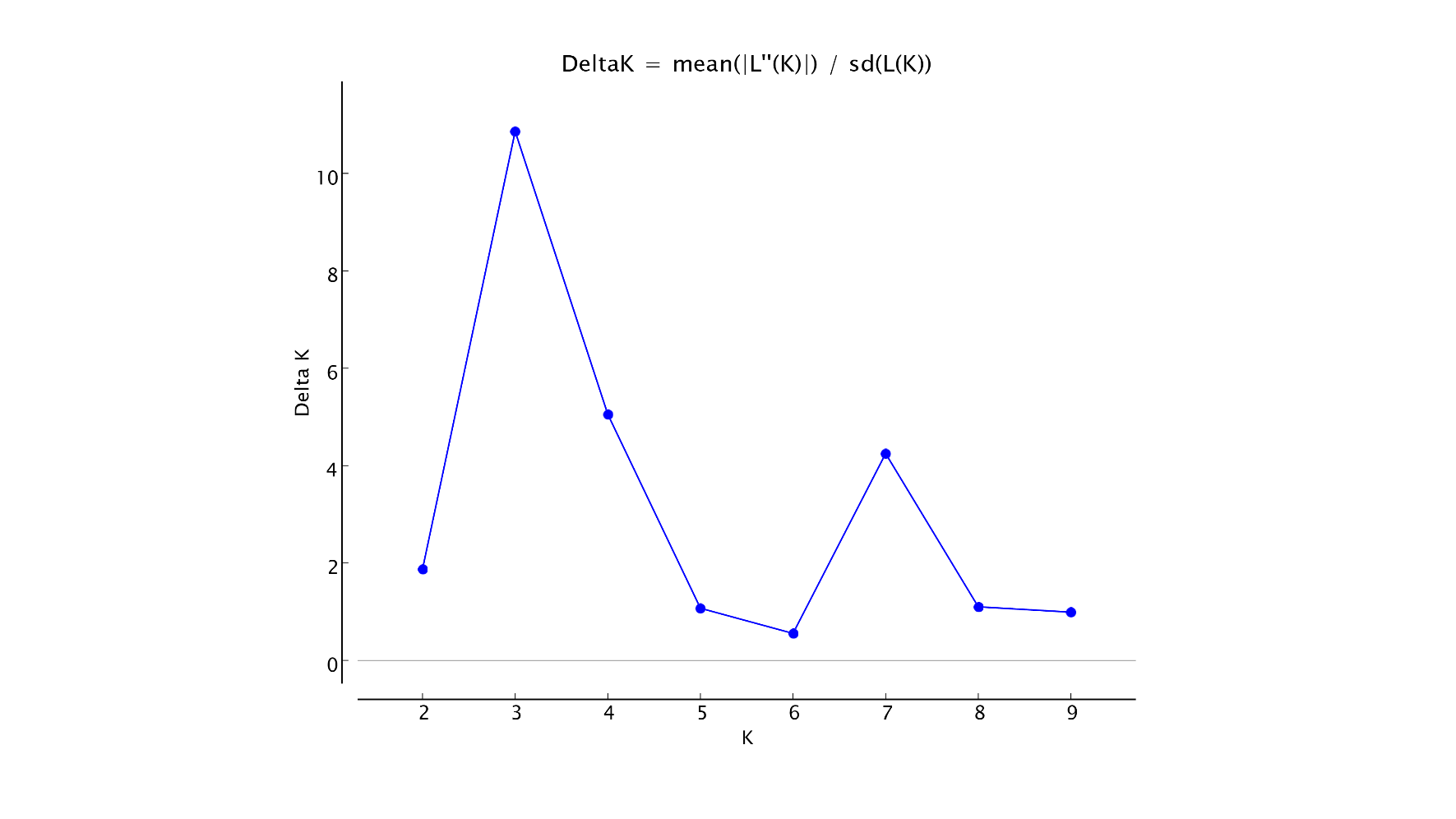

**Supplementary Fig. 1.** The most likely number of genetic clusters (K) within the full data set of 130 individuals based on the method described by Evanno et al. (2005). K values from ten separate runs, ranging from 1 to 10. The delta K graph identified the maximum value at K = 3.

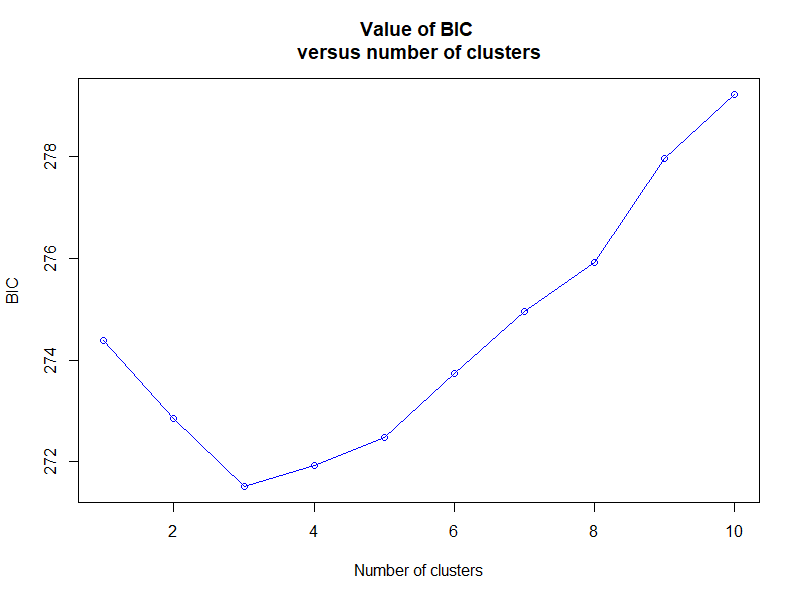

**Supplementary Fig. 2.** Bayesian information criterion (BIC) values for different numbers of clusters. The accepted true number of clusters was three.

**Supplementary Table 4.** Possible direct (first-degree) relationships of 42 IAC grapevine hybrids based on the maximum likelihood approach.

| **Offspring ID Number** | **Offspring Name** | **Candidate parent** | | **Pairs of loci compared** | | **Pairs of loci with mismatch** | | **Pair LOD score*** | |
| --- | --- | --- | --- | --- | --- | --- | --- | --- | --- |
| 9 | IAC 137-04 | | Sémillon | | 21 | | 1 | | 12.35 |
| 10 | IAC 138-22 (Máximo) | | Ravat 34 | | 19 | | 1 | | 10.50 |
| 11 | IAC 141-51 | | Sémillon | | 21 | | 0 | | 14.50 |
| 12 | IAC 158-12 | | Muscat Hamburg | | 16 | | 0 | | 8.49 |
| 14 | IAC 202-28 | | Muscat Hamburg | | 20 | | 0 | | 10.96 |
| 15 | IAC 202-43 | | Muscat Hamburg | | 21 | | 0 | | 11.70 |
| 19 | IAC 274-21 | | Ravat 34 | | 19 | | 1 | | 10.40 |
| 23 | IAC 338-04 | | *Vitis cinerea* | | 12 | | 0 | | 11.01 |
| 28 | IAC 387 | | Italia | | 21 | | 0 | | 11.90 |
| 30 | IAC 393-04 | | Muscat Hamburg | | 18 | | 0 | | 8.90 |
| 31 | IAC 393-05 | | Muscat Hamburg | | 21 | | 1 | | 12.20 |
| 32 | IAC 403-01 | | Sultanina | | 18 | | 1 | | 14.50 |
| 34 | IAC 408-01 | | *Vitis cinerea* | | 17 | | 0 | | 22.40 |
| 39 | IAC 486-03 | | Italia | | 20 | | 0 | | 11.80 |
| 52 | IAC 574-01 | | IAC 74-1 (Iara) | | 18 | | 0 | | 14.67 |
| 53 | IAC 583-03 | | Ruby Cabernet | | 14 | | 0 | | 13.26 |
| 54 | IAC 584-53 | | Sauvignon Gris | | 20 | | 0 | | 19.73 |
| 55 | IAC 589-02 | | Sémillon | | 18 | | 0 | | 12.05 |
| 56 | IAC 592-01 | | Ruby Cabernet | | 17 | | 0 | | 16.36 |
| 57 | IAC 594-03 | | Sémillon | | 20 | | 1 | | 15.32 |
| 61 | IAC 720-01 | | Carignane | | 16 | | 1 | | 11.49 |
| 63 | IAC 733-39 | | IAC 544-14 | | 16 | | 0 | | 9.74 |
| 69 | IAC 768-02 | | IAC 457-11 (Iracema) | | 21 | | 1 | | 8.31 |
| 70 | IAC 772-41 | | IAC 514-6 (Maria) | | 18 | | 0 | | 11.12 |
| 72 | IAC 778-04 | | IAC 544-14 | | 20 | | 0 | | 9.44 |
| 73 | IAC 804-13 | | IAC 583-03 | | 17 | | 0 | | 15.04 |
| 75 | IAC 822-21 | | IAC 405-06 | | 21 | | 0 | | 17.53 |
| 76 | IAC 842-04 (Eugenio) | | IAC 501-06 (Soraya) | | 15 | | 0 | | 13.25 |
| 77 | IAC 860-05 | | IAC 514-6 (Maria) | | 15 | | 0 | | 10.65 |
| 86 | IAC 901-01 | | IAC 457-11(Iracema) | | 20 | | 1 | | 6.65 |
| 88 | IAC 903-47 | | IAC 457-11(Iracema) | | 21 | | 1 | | 12.97 |
| 89 | IAC 904-11 | | IAC 514-6 (Maria) | | 16 | | 0 | | 12.38 |
| 90 | IAC 904-30 | | IAC 544-14 | | 20 | | 0 | | 11.59 |
| 92 | IAC 904-47 | | IAC 514-6 (Maria) | | 20 | | 0 | | 11.97 |
| 95 | IAC 915-02 | | IAC 457-11 (Iracema) | | 21 | | 1 | | 6.52 |
| 96 | IAC 918-52 | | IAC 457-11 (Iracema) | | 21 | | 1 | | 6.52 |
| 97 | IAC 960-11 | | IAC 138-22 (Máximo) | | 20 | | 1 | | 11.47 |
| 98 | IAC 960-12 | | Pinot Noir | | 15 | | 0 | | 12.14 |
| 101 | IAC 1025-17 | | IAC 772-41 | | 19 | | 0 | | 18.60 |
| 108 | IAC 1410-08 (Ezequiel) | | IAC 501-06 (Soraya) | | 21 | | 0 | | 23.42 |
| 113 | IAC 1726-03 (Roberta) | | IAC 871-18 | | 18 | | 0 | | 15.60 |
| 120 | Jd 930 (Moscatel de Jundiai) | | Seyve Villard 5276 | | 20 | | 1 | | 13.90 |

* A maximum of only one locus mismatch was allowed, and the parentage relationship was considered significant when the pair confidence probability was greater than 95% (LOD ≥ 6.40).

**Supplementary Table 5.** List of internal synonyms found in the grapevine hybrid collection of the Agronomic Institute of Campinas (IAC) by SSR analysis.

| **Group** | **IDs** | **Accession 1** | **Accession 2** | | **Proposed correct identification^a^** |
| --- | --- | --- | --- | --- | --- |
| 1 | 4-7 | IAC 31-01 | | IAC 82-01 | - |
| 2 | 28-39 | IAC 387 | | IAC 486-03 | - |
| 3 | 40-122 | IAC 496-15 | | SR 496-15 (Dr. Júlio) | SR 496-15 (Dr. Júlio) |
| 4 | 48-49 | IAC 544-14 | | IAC 547-02 | IAC 544-14 |
| 5 | 63-90 | IAC 733-39 | | IAC 904-30 | IAC 772-41 |
| 6 | 64-65 | IAC 740-01 | | IAC 746-03 | - |
| 7 | 36-71 | IAC 457-11 (Iracema) | | IAC 775-26 (Aurora) | IAC 457-11 (Iracema) |
| 8 | 80-81 | IAC 871-05 (Geni) | | IAC 871-13 (A Dona) | - |
| 9 | 95-96 | IAC 915-02 | | IAC 918-52 | IAC 915-02 |
| 10 | 8-99 | IAC 116-31 (Rainha) | | IAC 966-01 | IAC 116-31 (Rainha) |
| 11 | 115-116 | IAC 1848-04 | | IAC 1848-11 | - |
| 12 | 124-125 | SR 5010-08 | | SR 5010-21 | - |
| 13 | 127-130 | SR 5012-34 (Dona Emília) | | SR 507-08 | - |
| 14 | 126-128 | SR 501-17 (IAC Ribas) | | SR 501-33 | SR 501-17 (IAC Ribas) |

* Label correction based on the results of parentage and identity analysis.

**Supplementary Table 6.** Ampelographic characteristics of internal synonyms found in the IAC grapevine hybrid collection by SSR analysis.

| **ID** | **Genotype name** | **Synonymy group** | **OIV 225 Berry: color of skin** | **OIV 220 Berry: length** | **OIV 221 Berry: width** | **OIV 503 Berry: single-berry weight** | **OIV 202 Bunch: length (peduncle excluded)** | **OIV 203 Bunch: width** | **OIV 502 Bunch: single-bunch weight** | **OIV 505 Sugar content of must** | **Phenotypic variations on the upper surface of mature leaves** |
| --- | --- | --- | --- | --- | --- | --- | --- | --- | --- | --- | --- |
| 4 | IAC 31-01 | 1 | 2-rose | 5-medium | 5-medium | 3-low | 3-short | 1-very narrow | 1-very low | 5-medium | 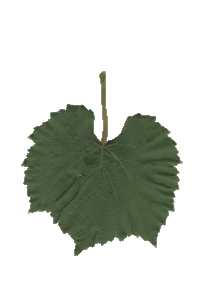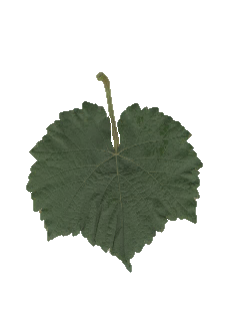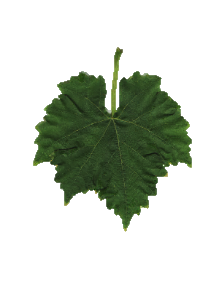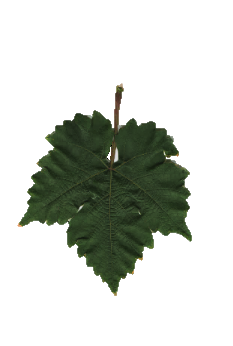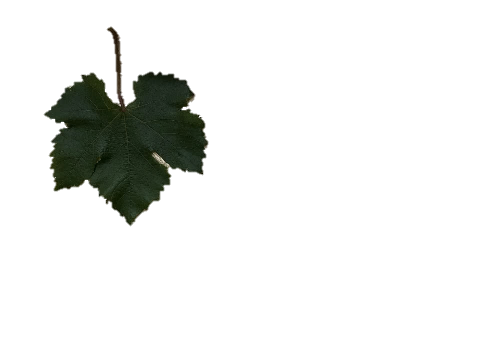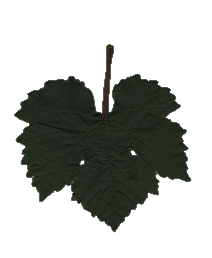 |
| 7 | IAC 82-01 | 1 | 2-rose | 5-medium | 5-medium | 3-low | 3-short | 1-very narrow | 1-very low | 5-medium |  |
| 28 | IAC 387 | 2 | 1-green–yellow | 5-medium | 5-medium | 3-low | 3-short | 1-very narrow or 3-narrow | 1-very low | 5-medium |  |
| 39 | IAC 486-03 | 2 | 1- green–yellow | 5-medium | 5-medium | 3-low | 3-short | 1-very narrow or 3-narrow | 1-very low | 5-medium |  |
| 40 | IAC 496-15 | 3 | 1- green–yellow | 3-short | 3-narrow | 1-very low | 3-short | 1-very narrow or 3-narrow | 1-very low | 7-high |  |
| 122 | SR 496-15 (Dr. Júlio) | 3 | 1- green–yellow | 3-short | 3-narrow | 1-very low | 3-short | 1-very narrow | 1-very low | 7-high |  |
| **ID** | **Genotype name** | **Synonymy group** | **OIV 225 Berry: color of skin** | **OIV 220 Berry: length** | **OIV 221 Berry: width** | **OIV 503 Berry: single-berry weight** | **OIV 202 Bunch: length (peduncle excluded)** | **OIV 203 Bunch: width** | **OIV 502 Bunch: single-bunch weight** | **OIV 505 Sugar content of must** | **Phenotypic variations on the upper surface of mature leaves** |
| 48 | IAC 544-14 | 4 | 6-blue–black | 3-short | 1-very narrow or 3-narrow | 1-very low | 3-short or 5-medium | 1-very narrow or 3-narrow | 1-very low | 5-medium or 7-high | 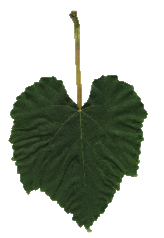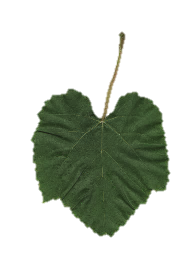 |
| 49 | IAC 547-02 | 4 | 6- blue–black | 3-short | 1-very narrow or 3-narrow | 1-very low | 3-short or 5-medium | 1-very narrow or 3-narrow | 1-very low | 5-medium or 7-high |  |
| 63 | IAC 733-39 | 5 | 2-rose | 3-short | 3-narrow | 1-very low | 1-very short or 3-short | 1-very narrow | 1-very low | 5-medium | 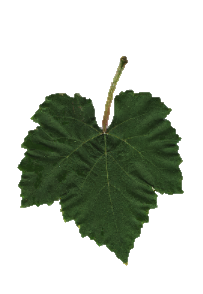 |
| 90 | IAC 904-30 | 5 | 2-rose | 3-short | 3-narrow | 1-very low | 1-very short or 3-short | 1-very narrow or 3-narrow | 1-very low | 5-medium | 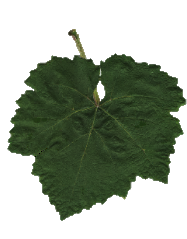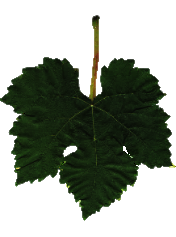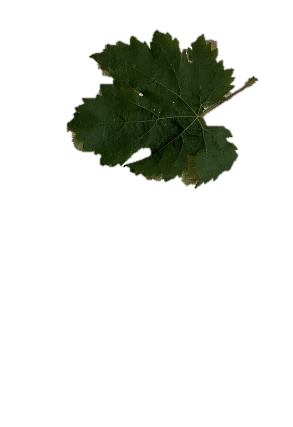 |
| 64 | IAC 740-01 | 6 | 6- blue–black | 5-medium | 5-medium | 3-low | 3-short | 3-narrow | 1-very low or 3-low | 5-medium |  |
| 65 | IAC 746-03 | 6 | 6- blue–black | 5-medium | 5-medium | 3-low | 3-short | 3-narrow | 1-very low or 3-low | 5-medium |  |
| **ID** | **Genotype name** | **Synonymy Group** | **OIV 225 Berry: color of skin** | **OIV 220 Berry: length** | **OIV 221 Berry: width** | **OIV 503 Berry: single berry weight** | **OIV 202 Bunch: length (peduncle excluded)** | **OIV 203 Bunch: width** | **OIV 502 Bunch: single bunch weight** | **OIV 505 Sugar content of must** | **Phenotypic variations on the upper surface of mature leaves** |
| 36 | IAC 457-11 (Iracema) | 7 | 1-green–yellow | 3-short or-5- medium | 3-narrow | 1-very low or-3-low | 1-very short or 3-short | 1-very narrow | 1-very low | 5-medium or 7-high | 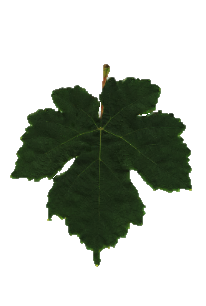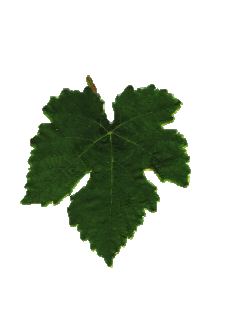 |
| 71 | IAC 775-26 (Aurora) | 7 | 1-green–yellow | 3-short | 3-narrow | 1-very low | 1-very short or 3-short | 1-very narrow | 1-very low | 5-medium |  |
| 80 | IAC 871-05 (Geni) | 8 | 2-rose | 5-medium | 3-narrow | 1-very low or-3-low | 3-short or 5-medium | 1-very narrow or 3-narrow | 1-very low | 5-medium | 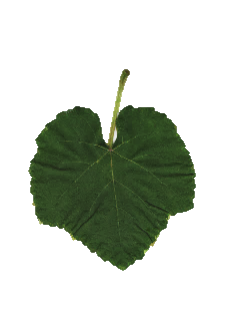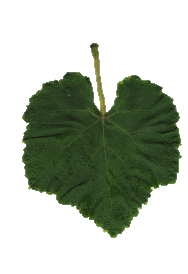 |
| 81 | IAC 871-13 (A Dona) | 8 | 2-rose | 5-medium | 3-narrow | 1-very low or-3-low | 3-short or 5-medium | 1-very narrow or 3-narrow | 1-very low | 5-medium |  |
| 95 | IAC 915-02 | 9 | 6-blue–black | 3-short | 3-narrow | 1-very low | 5-medium | 1-very narrow or 3-narrow | 1-very low | 5-medium or 7-high | 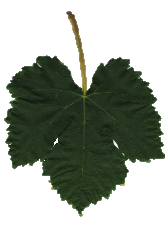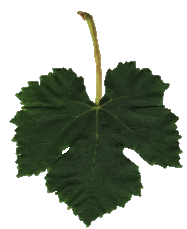 |
| 96 | IAC 918-52 | 9 | 6-blue–black | 3-short | 3-narrow | 1-very low | 5-medium | 3-narrow | 1-very low | 5-medium or 7-high |  |

| **ID** | **Genotype name** | **Synonymy group** | **OIV 225 Berry: color of skin** | **OIV 220 Berry: length** | **OIV 221 Berry: width** | **OIV 503 Berry: single-berry weight** | **OIV 202 Bunch: length (peduncle excluded)** | **OIV 203 Bunch: width** | **OIV 502 Bunch: single-bunch weight** | **OIV 505 Sugar content of must** | **Phenotypic variations on the upper surface of mature leaves** |
| --- | --- | --- | --- | --- | --- | --- | --- | --- | --- | --- | --- |
| 8 | IAC 116-31 (Rainha) | 10 | 1-green–yellow | 3-short | 3-narrow | 1-very low | 3-short | 1-very narrow | 1-very low | 7-high | 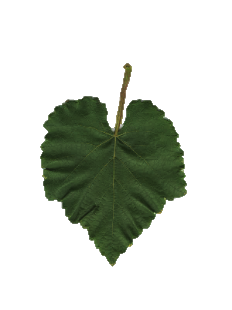 |
| 99 | IAC 966-01 | 10 | 1-green–yellow | 3-short | 3-narrow | 1-very low | 3-short | 1-very narrow or 3-narrow | 1-very low | 7-high | 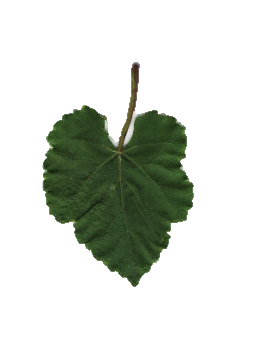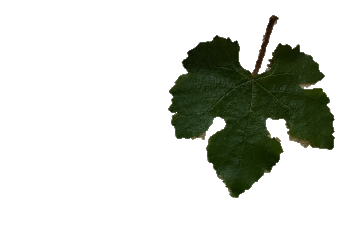 |
| 115 | IAC 1848-04 | 11 | - | - | - | - | - | - | - | - |  |
| 116 | IAC 1848-11 | 11 | - | - | - | - | - | - | - | - | 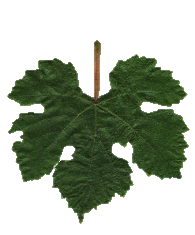 |
| 124 | SR 5010-08 | 12 | 6-blue–black | 3-short | 3-narrow | 1-very low | 5-medium | 3-narrow | 1-very low | 5-medium | 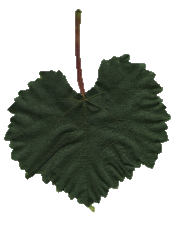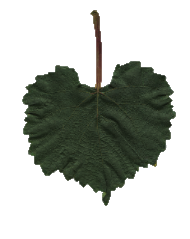 |
| 125 | SR 5010-21 | 12 | 6-blue–black | 3-short | 3-narrow | 1-very low | 5-medium | 3-narrow | 1-very low | 5-medium |  |

| **ID** | **Genotype name** | **Synonymy group** | **OIV 225 Berry: color of skin** | **OIV 220 Berry: length** | **OIV 221 Berry: width** | **OIV 503 Berry: single-berry weight** | **OIV 202 Bunch: length (peduncle excluded)** | **OIV 203 Bunch: width** | **OIV 502 Bunch: single-bunch weight** | **OIV 505 Sugar content of must** | **Phenotypic variations on the upper surface of mature leaves** |
| --- | --- | --- | --- | --- | --- | --- | --- | --- | --- | --- | --- |
| 127 | SR 5012-34 (Dona Emília) | 13 | 6-blue–black | 3-short | 3-narrow | 1-very low | 1-very short | 1-very narrow | 1-very low | 5-medium | 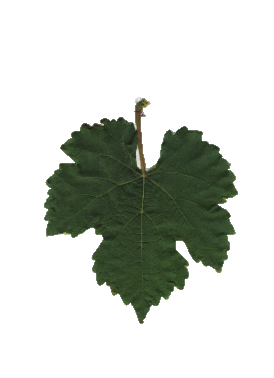 |
| 130 | SR 507-08 | 13 | 6-blue–black | 3-short | 3-narrow | 1-very low | 1-very short | 1-very narrow | 1-very low | 5-medium | 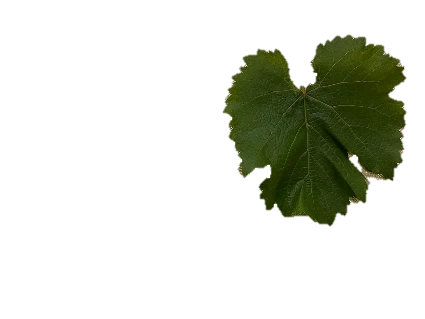 |
| 126 | SR 501-17 (IAC Ribas) | 14 | 1-green–yellow | 3-short or-5- medium | 3-narrow or-5- medium | 1-very low or-3-low | 3-short | 1-very narrow or 3-narrow | 1-very low | 5-medium | 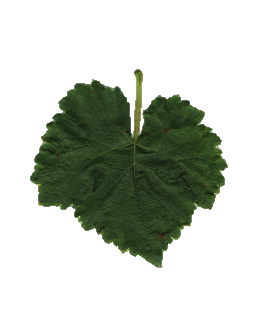 |
| 128 | SR 501-33 | 14 | 1-green–yellow | 3-short or-5-medium | 3-narrow or-5-medium | 1-very low or-3-low | 1-very short or 3-short | 1-very narrow or 3-narrow | 1-very low | 5-medium | 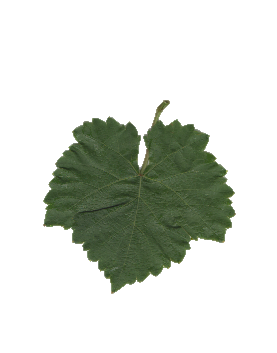 |
